# Geospatially informed representation of spatial genomics data with SpatialFeatureExperiment

**DOI:** 10.1101/2025.02.24.640007

**Authors:** Lambda Moses, Alik Huseynov, Joseph M Rich, Lior Pachter

## Abstract

SpatialFeatureExperiment is a Bioconductor package that leverages the versatility of Simple Features for spatial data analysis and SpatialExperiment for single-cell -omics to provide an expansive and convenient S4 class for working with spatial -omics data.

SpatialFeatureExperiment can be used to store and analyze a variety of spatial -omics data types, including data from the Visium, Xenium, MERFISH, SeqFish, and Slide-seq platforms, bringing spatial operations to the SingleCellExperiment ecosystem.

## Main

Single cell, nucleus, and spatial -omics technologies have become standard and essential tools for a large swath of molecular biology, where they are used to understand tissue heterogeneity and biological functions^1^. Data structures for storing and analyzing the data generated by these technologies are foundational; they can enable and facilitate complex queries of the data, or, if not well designed, can limit the scope of analyses. Novel data structures underlying single cell data analysis include Seurat^2^, SingleCellExperiment^3^, and AnnData^4^. These data structures organize the data as gene count matrices along with cell and gene metadata, and they are also used to manage results of common analyses such as clustering and dimension reduction. The data structures are distributed with tools for access and manipulation; specifically operations for tasks such as subsetting, combining multiple objects, and retrieving and setting fields of the structure are implemented to simplify bookkeeping.

Data structures for spatial -omics data analysis have largely resulted from modifications or extensions of existing non-spatial single-cell/nucleus -omics data structures. However, the spatial information necessitates data representation and operations that are not relevant for non-spatial data such as those arising from single-cell RNA-seq or single-nucleus RNA-seq experiments. Due to the importance of data structures, there have been several parallel efforts to enhance single cell data structures for spatial data, such as new spatial fields implemented for objects in Seurat, SpatialExperiment^5^, Giotto^6^, Staffli^7^, SpatialData^8^, and MoleculeExperiment^9^. These data structures all store spatial coordinates but have varying support for different types of spatial information and spatial data operations (Supplementary Tables 1-2). Some of the data structures are less generalizable to different spatial -omics technologies; Seurat has very different internal structures for sequencing based and imaging based technologies, and some functionalities of SpatialExperiment^10^ and Staffli are specific to Visium. Currently the R interface to Python’s SpatialData provides reader as well as plotting functions and stores data objects in Zarr format^11^.

While the development of spatial analysis within -omics is relatively new, spatial data structures are well-established in geography. There are two main types of spatial data, which are present in both geography and spatial -omics. The first type is vector, represented by numerical coordinates of points and vertices of lines and polygons. In geography, these can be locations of landmarks such as landmarks (points), roads and railway (linestrings), and parks (polygons) (Figure 1A). In spatial -omics, these can be cell centroids (points), transcript spot locations (points), cell segmentation polygons, tissue boundaries (polygons), and pathologist annotated histological regions (polygons) (Figure 1A). Emerging from needs in geography, Simple Features^12^ is a standardized open format to represent vector data. Spatial operations on Simple Features such as finding spatial intersections and performing spatial buffering (Supplementary Figure 2), as well as input/output (IO) of various vector file formats, are implemented in well-established C++ libraries such as GDAL and GEOS, which can be accessed from the sf R package. The second type of spatial data in geography is raster, i.e. image-like arrays with data at each pixel, registered to a spatial coordinate system. In geography, raster data often comes from remote sensing, such as satellite images in Google Maps (Figure 1A). In spatial -omics, histology images and cell segmentation masks constitute raster data (Figure 1A). GDAL supports raster data IO for a variety of formats, including chunked formats such as NetCDF and Zarr. In R, terra^13^ and stars^14^ are the main packages for raster data operations.

**Figure 1:**
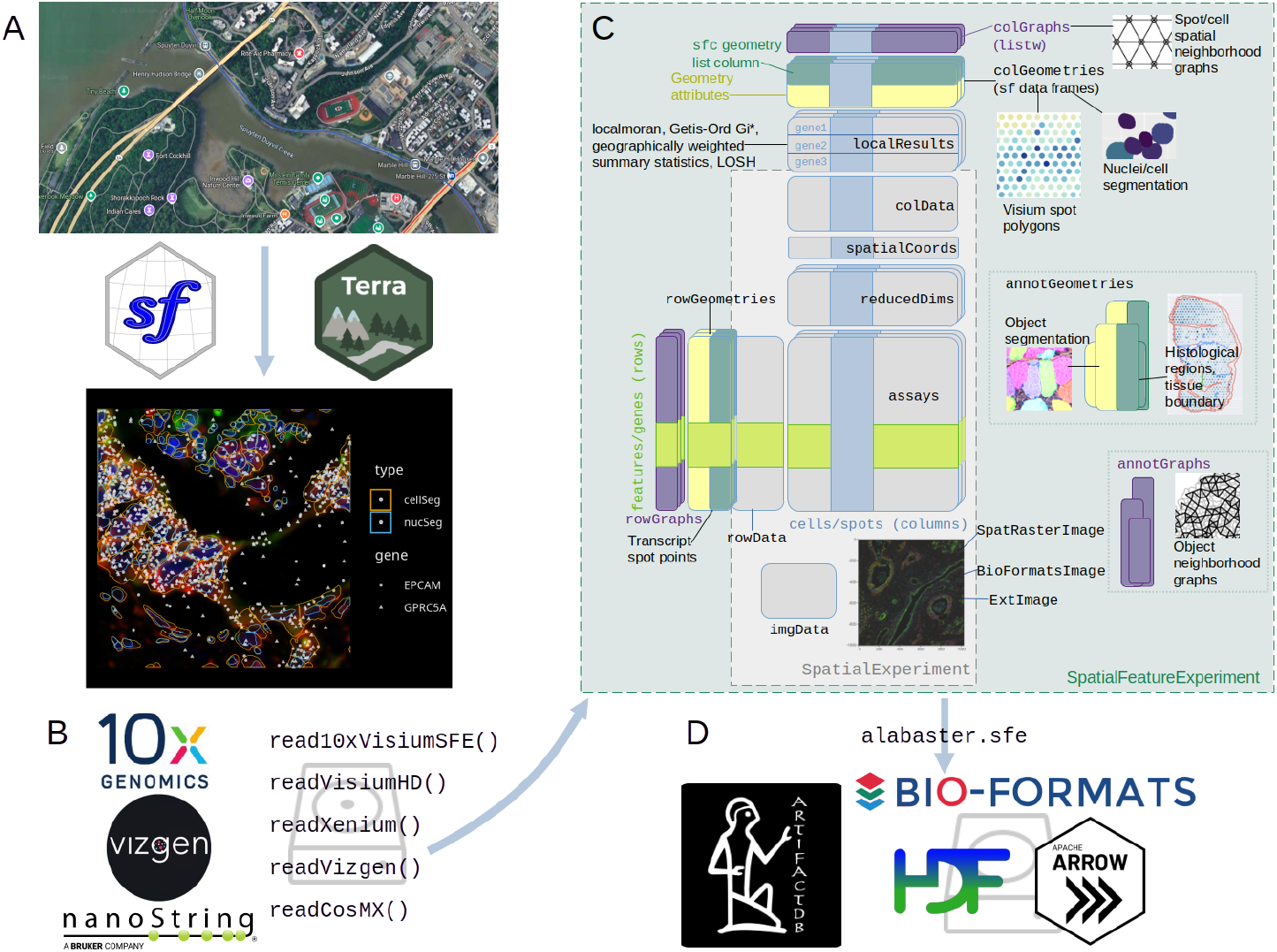
Schematic of the SpatialFeatureExperiment project. A) Geospatial tools such as sf and terra make the SFE object like a multi-layered map with vector and raster data aligned in a common coordinate system. B) SFE implements read functions to conveniently read output from commercial spatial transcriptomics technologies into R. C) Schematic of the SFE object. D) The alabaster.sfe package builds upon ArtifactDB and uses BioFormats, HDF5, and Apache Arrow for out of memory representation of SFE data.

To take advantage of the opportunity to leverage the tools of Spatial Features in the spatial -omics setting, we developed a framework that can also take advantage of expansive R toolkits for single-cell -omics. SingleCellExperiment (SCE) is a well-established and stable data structure in the R programming language for non-spatial single cell -omics data on Bioconductor, as part of an ecosystem of packages for preprocessing and analyses of single cell -omics data. SpatialExperiment (SPE) extends SCE by adding fields to store cell centroid coordinates and images.

We have developed SpatialFeatureExperiment (SFE), which extends SPE by using sf for vector geometries beyond centroids, and terra and EBImage^15^ for raster images, allowing for spatial operations on the geometries and images. Extending SCE and SPE, SFE is seamlessly integrated into the existing ecosystem of spatial and non-spatial tools, such as scater^16^ and scran^17^ for basic non-spatial preprocessing, plotting, and analyses, and BayesSpace^18^ and lisaClust^19^ for spatial analyses. In addition, inheriting from SCE, SFE can facilitate DelayedArray^20^ to work with large datasets out of memory, which are increasing common due to the increasing number of cells in imaging based spatial -omics data^1^ and development of technologies with increasing resolution such as Stereo-seq^21^ and Visium HD^22^.

With geometries and images registered to a spatial coordinate system, an SFE object can be thought of as a multi-layered map, just like Google Maps with vector site locations, rivers, roads, and city boundaries on top of raster satellite images (Figure 1A). SFE is designed to be generalizable to any spatial -omics technology where a feature count matrix is central to analysis. Functions to read output from Visium, VisiumHD, Vizgen MERFISH, Xenium, and CosMX have been implemented and more commercial technologies such as Molecular Cartography and Curio Seeker will be supported in the near future (Figure 1B). SFE has a website documenting all user-facing functions and showcasing operations of SFE objects in vignettes.

Vector geometries in SFE can be associated with dimensions of the gene count matrix, as a special case of gene (rowGeometries) or cell (colGeometries) metadata (Figure 1C). For Visium, colGeometry can be spot centroids or spot polygons with diameter from the Space Ranger output. For imaging based technologies such as MERFISH and Xenium, colGeometry can be cell and nucleus segmentation polygons, and rowGeometry transcript spots, including those outside cells. In addition, there can be geometries associated with the dataset that don’t correspond to dimensions of the gene count matrix, such as tissue boundaries, histological regions, and cell segmentation in Visium; these are stored in annotGeometries (Figure 1C).

SFE currently implements three different image classes to leverage the advantages of different image processing traditions (Figure 1A). In all image classes, a spatial extent to register the image to the spatial coordinate system is required. The SpatRasterImage class is based on terra, from the geospatial tradition; support of the extent is built-in and vector geometries can be used to extract data from images for multi-modal analyses and quality control. Here along with total transcript counts, image pixel values are used to identify low quality spots at the edge of the tissue (Figure 2F). For images that don’t fit into memory, only the metadata is read and operations such as flipping and transposing can be performed chunk wise, never loading the entire image into memory as implemented in terra. Large images can also be efficiently downsampled for plotting with terra. GDAL is used to read a wide variety of raster formats, potentially facilitating interoperability with the Zarr-based SpatialData format. The ExtImage class complements SpatRasterImage with image processing functionalities from the EBImage^15^ package, such as Otsu thresholding, Haralick feature extraction, watershed segmentation, and morphological operations, although the image is loaded into memory. The BioFormatsImage class represents images without reading them into memory, and determines spatial extent from metadata from OME-TIFF files and potentially other formats supported by BioFormats (Figure 1D). Affine transformations are stored in the metadata as well. Conversion between classes is implemented, such as to load the image of BioFormatsImage into memory as ExtImage, or to convert ExtImage into SpatRasterImage for vector-raster interaction. All 3 classes inherit from a virtual class AlignedSpatialImage (Methods), which can be used to implement new classes to extend SFE.

**Figure 2:**
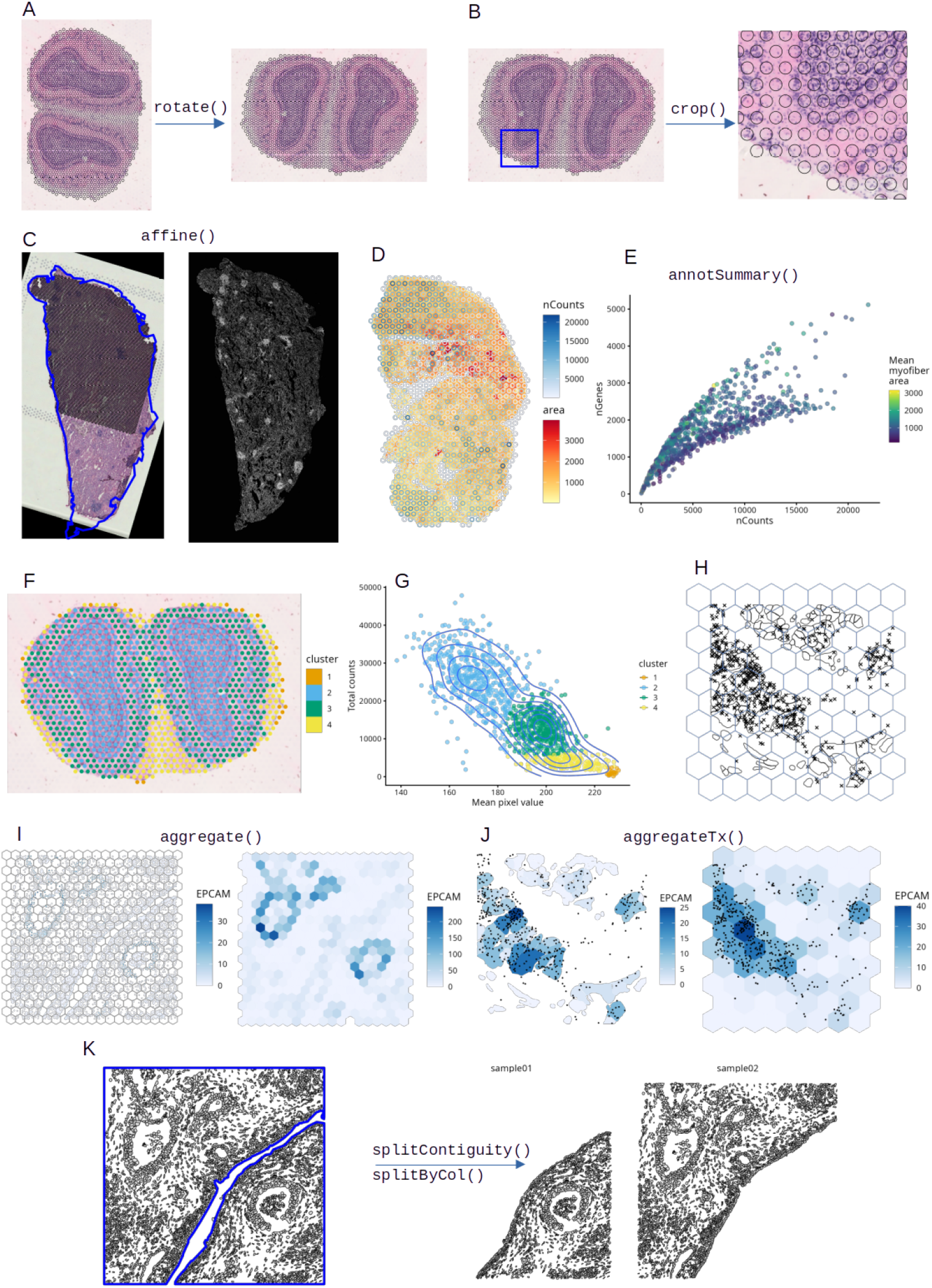
Summary of spatial operations implemented in SFE. A) The SFE object, with geometries and images aligned, can be rotated into a canonical orientation. B) Cropping the SFE object. C) General affine transformation can be used to align serial sections profiled with different technologies; here a Visium dataset (H&E image and spots shown as black circles) is registered to Xenium (fluorescent image and blue outline on H&E). D) Geometric predicates can be used to relate annotGeometries (myofiber polygons) to colGeometries (Visium spots). E) Scatter plot of number of genes detected per Visium spot (nGenes) vs. total transcript count (nCounts), colored by mean area of myofibers that intersect the spot, showing two branches; larger myofibers tend to have lower nCounts for the same nGenes. F) Mouse olfactory bulb Visium spots colored by clusters based on nCounts and pixel intensity from the H&E image. G) Scatter plot of nCounts vs. H&E pixel intensity, colored by clusters. Point density is shown as contours. H) A subset of a Xenium dataset from human pancreas showing transcript spots of the gene EPCAM plotted as points, cell segmentation polygons, and hexagonal bins. I) Cell-level data can be aggregated by spatial bins according to which bin the cell is located in; here cell-level counts of EPCAM are summed over bins. J) Transcripts can be assigned to any spatial region they belong to to generate new gene count matrices, such as cell segmentation polygons and hexagonal bins. K) Tissue boundary regions are plotted in blue lines, and cell segmentations are plotted in orange. The SFE object can be split with tissue region annotations.

Affine transformation can be applied to the entire SFE object including all the geometries and images like a stack of maps; this is useful to transform the data into a canonical orientation (Figure 2A) or to align it to data from a different modality such as aligning Visium and Xenium data from serial sections (Figure 2C). The transformations can be named – such as mirroring, transposing, translation, and scaling – or can be specified with a transformation matrix (Supplementary Table 3). Here all geometries and images are transformed together while remaining aligned akin to a multi-layered map.

Spatial coordinates of transcript spots are available from smFISH-based technologies; subcellular transcript localization can be of biological interest^23^. These can be optionally read with functions that read output from commercial smFISH-based technologies, which reformat the transcript spots into Simple Features written to disk in GeoParquet format for faster reading for future use (Figure 1D). The reformatting can also be performed independently from the read functions.

With sf, the geometries can be used to perform operations on the SFE object. These include spatial cropping (Figure 2B) with a bounding box or any geometry specifying a region of interest. Spatial predicates can relate data associated with different geometries. For example, in an injured mouse skeletal muscle Visium dataset^24^, smaller and more tightly packed myofibers tend to be in Visium spots with higher total transcript counts (Figure 2D-E). This confirms the finding that in spatial transcriptomics, library size is biologically relevant and should not be treated as a technical artifact as is commonly done for scRNA-seq^25^. Geometries can be used to aggregate data at the cell (Figure 2I) and transcript spot (Figure 2J) levels. The implementation is flexible, allowing for aggregation by any geometry including cell segmentation polygons and regular grids. When aggregating cellular level data, any summary function can be used, not limited to standard functions such as sum and mean. Conversely, geometries can be used to split an SFE object (Figure 2K). For example, with an annotation of histological regions, cells in different regions can be assigned to different SFE objects for further analyses.

To facilitate single-cell analyses and simplify bookkeeping, Seurat and SCE objects store analysis results such as cell cluster labels and cell projections in reduced dimensional space within the object (Figure 1C). Similarly, the SFE object has fields to store exploratory spatial data analysis (ESDA) results from Voyager^26^. Global univariate statistics, e.g. Moran’s I and Geary’s C that yield one result for an entire dataset, are stored in rowData, and local univariate statistics, e.g. local Moran’s I, that return results at each location, as well as bivariate statistics such as local Lee’s L for pairs of genes are stored in the new field localResults analogous to reducedDims for each gene separately (Figure 1C). When the rows or columns of the SFE object are selected, the corresponding ESDA results are filtered as well. The ESDA results can then be retrieved from these fields for visualization and secondary analysis such as clustering by local Moran’s I, a continuous analogue of the concordex spatial clustering method^27^ (Supplementary Figure 1C).

Because many ESDA methods require a spatial neighborhood graph, SFE supports many methods to find spatial neighborhood graphs such as triangulation and methods to prune the triangulated graphs, k nearest neighbors, distance-based neighbors, and polygon contiguity – all can be called with a uniform user interface (Supplementary Table 4). The spatial neighborhood graphs are stored in the colGraph and annotGraph fields for colGeometries and annotGeometries respectively, allowing for ESDA for attributes of annotGeometries such as area of myofibers in Visium dataset as well as colGeometries such as Visium spots.

To manage the growing size of the data, SFE allows out of memory components with terra and BioFormatsImage for images and DelayedArray for gene count matrices. The alabaster.sfe package implements language-agnostic on-disk serialization of SFE objects to facilitate data sharing and interoperability between languages. The SPE part of SFE is serialized with existing alabaster packages developed under ArtifactDB^28^, and the geometries are saved as GeoParquet which is designed for efficiency and interoperability.

SpatialFeatureExperiment extends SingleCellExperiment and SpatialExperiment with sf and additional image classes incorporating the geospatial raster, image processing, and microscopy traditions. The geometries can interact with each other and with raster images to relate data from different modalities. These new functionalities allow users to place the spatial information front and center and think about the data as a map in addition to gene count matrices, making new spatial insights possible. Because SFE inherits from SPE and SCE, which in turn inherits from SummarizedExperiment, non-spatial operations such as matrix-style subsetting and getter and setter functions for non-geometry fields are inherited. These include imgData from SPE for images, reducedDims from SCE for dimension reduction results, and colData and rowData from SummarizedExperiment for column and row metadata for the gene count matrix. The new fields for geometries and spatial results have getter and setter functions that conform to SCE conventions. ColGeometries, rowGeometries, annotGeometries, colGraphs, rowGraphs, and annotGraphs getters and setters have similar user interface to reducedDims getters and setters. As part of the SCE ecosystem, SFE may be adapted to integrate into tidyOmics^29^.

With increasing amount of geospatial data, interoperable database frameworks such as DuckDB, TileDB, and Apache Sedona have been developed to manage and operate on out-of-memory spatial data, with GDAL integration. The SFE framework can be extended to incorporate these database solutions to accommodate the increasing size of spatial -omics data (Methods).

## Code availability

SpatialFeatureExperiment is freely available from Bioconductor at https://www.bioconductor.org/packages/release/bioc/html/SpatialFeatureExperiment.html

SFE’s documentation website is available at https://pachterlab.github.io/SpatialFeatureExperiment/ alabaster.sfe has been submitted to Bioconductor and is currently under review. It is freely available at https://github.com/pachterlab/alabaster.sfe

Code to reproduce figures is available at https://github.com/pachterlab/MHRP_2024

## Acknowledgement

We thank SpatialExperiment author Dario Righelli for suggestions in the early development of SFE. We thank the Bioconductor community whose work and infrastructure SFE is based on and who provided feedback on this package. We thank Andrew Ghazi and Eva Freckmann for contributing to SFE’s codebase, Aaron Lun for his support during the development of alabaster.sfe, Alex Mahmoud for building the Bioconductor Docker images and Galaxy instance with which SFE workshops have been held, and Vincent Carey for reviewing and contributing to the alabaster.sfe package. This work was supported in part by the Icelandic Research Fund Project [218111-051], NIH 5UM1HG012077-02, the Health + Life Science Alliance Heidelberg Mannheim MULTI-SPACE, and state funds approved by the State Parliament of Baden-Württemberg.

## Author contributions

LM and AH jointly implemented the SFE R package. LM conceived the idea of SFE, implemented alabaster.sfe, and drafted most of the manuscript, some of whose supplementary tables were made by AH. AH, JR, and LP reviewed and edited the manuscript.

