## Supplementary methods, figures, and tables for "Geospatially informed representation of spatial genomics data with SpatialFeatureExperiment"

### Online methods

#### Implementation of SFE

##### Relevant internals of SPE and SCE

The SPE class extends SCE by adding a new `spatialCoords` field to store cell or Visium spot centroid coordinates, and an `imgData` field for the images and their metadata (Figure 1C). Internally, `spatialCoords` is a matrix whose rows correspond to cells and whose columns correspond to spatial dimensions; it's stored in an internal field in the SCE object, `int_colData`, as internal column metadata that should not be accessed directly by the user but can be accessed or modified by the developer-provided getter or setter function.

In SCE, reducedDims, which stores cell projections in dimension reduction spaces, uses `int_colData` internally. `reducedDims` allows for multiple projections, such as PCA, NMF, tSNE, and UMAP. int_colData is an S4 DataFrame implemented in the `S4Vectors` package from Bioconductor, which allows for great flexibility in the contents of the columns. A column can be an atomic vector of any type, and all columns in the `DataFrame` must have the same length, that is the number of rows of the `DataFrame`. A column can also be a list of the same length as the number of rows, or a matrix, S3 `data.frame`, or S4 `DataFrame` with any number of columns but the same number of rows as the `DataFrame`. The `int_colData DataFrame`, as column metadata, must have the same number of rows as the number of columns in the gene count matrix in the assays. `reducedDims` is a `DataFrame`, each column of which is a matrix for the cell projections in each dimension of a dimension reduction. Other information about the dimension, such as PCA gene loadings and variance explained, is stored as attributes of the matrix. In turn, the entire `reducedDims` is a column in `int_colData`, so is the `spatialCoords` matrix. All the cell projection matrices can be retrieved from the SCE object with `reducedDims(sce)` and set with `reducedDims(sce) <- value`. Individual cell projection matrices can be get with a string name or an integer index, such as `reducedDim(sce, 1)` or `reducedDim(sce, "PCA")`, and similarly set with `reducedDim(sce, "PCA") <- value`.

`imgData` does not correspond to dimensions of the gene count matrix in the assays; it's stored in internal metadata, or `int_metadata`. `imgData` is a `DataFrame` with one list column for the images themselves, and other columns for image ID, sample ID, and a scale factor that aligns the image to Visium spots based on the `scalefactors_json.json` file in Space Ranger output. In SPE, the sample ID, or sample_id, serves to distinguish between spots from different tissue sections, as different sections can have spots with overlapping spatial coordinate values. For the images, SPE implements the following new S4 classes:

1. `VirtualSpatialImage`: a virtual class allowing for new classes expending SPE. All image classes in SPE derivatives must inherit from this class.

2. `SpatialImage`: the image is loaded into memory as an array of hex colors.
3. `StoredSpatialImage`: the image is on disk and only the file path is in memory; the whole image will be read into memory when plotted.
4. `RemoteSpatialImage`: similar to `StoredSpatialImage`, but the image is online and the URL is in memory.

#### Internal column and row metadata

SFE extends SPE by adding the geometry and graph fields, `localResults`, and new image classes.

`colGeometries` are `sf` data frames (derived from S3 `data.frame`s with a geometry list column) associated with the entities that correspond to columns of the gene count matrix, such as Visium spots or cells. The geometries in the `sf` data frames can be Visium spot centroids, Visium spot polygons, or for datasets with single cell resolution, cell or nuclei segmentations. Multiple `colGeometries` can be stored in the same SFE object, such as one for cell segmentation and another for nuclei segmentation. There can be non-spatial, attribute columns in a `colGeometry` rather than `colData`, because the `sf` class allows users to specify how attributes relate to geometries, such as "constant", "aggregate", and "identity"[1]. Internally, each `colGeometry` `sf` data frame is a column in the `colGeometries DataFrame`, which in turn is a column in `int_colData` just like `reducedDims`. Getters and setters intentionally mimic those of `reducedDims`. So all `colGeometries` can be retrieved with `colGeometries(sfe)`, or set with `colGeometries(sfe) <- value`. Individual `colGeometry` can be retrieved with either a string name or a integer index such as `colGeometry(sfe, "spotPoly")` or `colGeometry(sfe, 1)`, and similarly set with `colGeometry(sfe, "spotPoly") <- value`. However, there's another argument sample_id to get or set colGeometries for one sample specifically when there are multiple samples in the SFE object for joint non-spatial SCE analyses.

While we use terra for images in SFE and terra also has vector geometry functionalities, sf is used for vector geometries here to utilize its larger ecosystem of packages, including ggplot2[2], sfheaders[3], lwgeom[4], rmapshaper[5], sfarrow[6], and smoothr[7], as well as sf's Tidyverse integration which is very useful for QC and downstream analyses using the geometries. Using sf also has a historical reason, that the SFE idea originated from colGeometries, and image support was added later. In a future implementation, in order to support out of memory geometric representation and operation such as with geoarrow[8] and DuckDB, a more expansive class such as sdf[9] may be supported. This should not be too challenging thanks to DataFrame's flexibility in the content of columns.

An SFE object can have any number of colGeometries with any names and geometry types. For example, for smFISH-based data, the SFE object can have cell segmentation, nucleus segmentation, cell segmentations from different algorithms, cell or nucleus centroids, and so on. When a cell is detected by membrane stain but a nucleus is not detected, then the corresponding nucleus segmentation can be an empty geometry. Cell segmentation is usually of type POLYGON, but nucleus segmentation can be MULTIPOLYGON when some cells have multiple nuclei. Cell centroids are of type POINT.

rowGeometries are similar to colGeometries, but support entities that correspond to rows of the gene count matrix, such as genes. It is currently used for transcript spots in smFISH-based data; transcript spots – including those outside cells – are represented as a MULTIPOINT geometry for each gene. Internally, analogous to int_colData, internal row metadata in SCE is in int_elementMetadata. rowGeometries are stored in int_elementMetadata. rowGeometries can be sample-specific, in which case the name of the sample the rowGeometry is for is appended to the name of the rowGeometry and the rowGeometry getter and setter take the sample name into account.

localResults are similar to reducedDims in SingleCellExperiment, but stores results from univariate and bivariate local spatial analysis results, such as from localmoran, Getis-Ord Gi*, and local spatial heteroscedasticity (LOSH). Unlike in reducedDims, for each type of results (type is the type of analysis such as Getis-Ord Gi*), each feature (e.g. gene) or pair of features for which the analysis is performed has its own results. The local spatial analyses can also be performed for attributes of colGeometries and annotGeometries in addition to gene expression and colData. Results of multivariate spatial analysis such as MULTISPATI PCA[10] can be stored in reducedDims.

Using int_colData and int_elementMetadata in SCE simplifies bookkeeping since SCE maintains the match between rows of the internal col and row metadata and dimensions of the gene count matrix when the SFE object is subsetted or concatenated.

#### Internal metadata

colGraphs are spatial neighborhood graphs of cells or spots. The graphs have class listw (spdep package), and the colPairs field of SCE was not used so no conversion is necessary to use the numerous spatial dependency functions from spdep[11], such as those for Moran's I, Geary's C, Getis-Ord Gi*, LOSH, etc. Conversion is also not needed for other classical spatial statistics packages such as spatialreg[11] and adespatial[12]. The graphs are stored in the SFE object so they can be reused for different analysis tasks. Any number of colGraphs can be present, such as k nearest neighbors, distance-based neighbors, triangulation, and polygon contiguity. The SFE function findSpatialNeighbors gives a uniform user interface to many spatial neighborhood graph functions in spdep, but uses the much faster and flexible implementations in BiocNeighbors[13] instead for k nearest neighbor and distance-based graphs.

rowGraphs are similar to colGraphs. A potential use case may be spatial colocalization of transcripts of different genes. However, while it’s implemented, it has not been used.

annotGeometries are sf data frames associated with the dataset but not directly with the gene count matrix, such as tissue boundaries, histological regions, cell or nuclei segmentation in Visium datasets, and etc. These geometries are stored in this object to facilitate plotting and using sf for operations such as to find the number of nuclei in each Visium spot and which histological regions each Visium spot intersects. Unlike colGeometries and rowGeometries, the number of rows in the sf data frames in annotGeometries is not constrained by the dimension of the gene count matrix and can be arbitrary. Internally, annotGeometries form a named list in int_metadata.

annotGraphs are similar to colGraphs and rowGraphs, but are for entities not directly associated with the gene count matrix, such as spatial neighborhood graphs for nuclei in Visium datasets, or other objects like myofibers. These graphs are relevant to spdep analyses of attributes of these geometries such as spatial autocorrelation in morphological metrics of myofibers and nuclei. With geometry operations with sf, these attributes and results of analyses of these attributes (e.g. spatial regions defined by the attributes) may be related back to gene expression.

Internally, the spatial graphs are in a DataFrame whose columns correspond to sample_ids and whose rows correspond to margins (row, column, and annotation). Each element in the data frame is a named list of all the spatial graphs for the sample and margin of interest. Getter and setter (e.g. colGraphs, colGraph, annotGraph) have the same arguments as the geometry getters and setters.

#### Feature data

Besides gene expression, spatial analyses can be performed on columns of colData such as cell-level QC metrics, non-geometry columns of colGeometry and annotGeometry, and cell projections in reduced dimensional spaces (for example Supplemenatry Figure 1B). Results of global spatial analyses are stored in feature data (Supplementary Figure 1A). For colData, which is of class DataFrame, feature data is stored in mcols as implemented in S4Vectors. For colGeometry, annotGeometry, and reducedDim, the results are stored in the attributes. The results can be retrieved with getters colFeatureData(), geometryFeatureData(), and reducedDimFeatureData(). When these are subsetted by column, the feature data corresponding to the removed columns are also removed.

#### SFE image classes

In SPE, the images are only used for visualization but not for analyses and the whole image is read into memory when necessary. SFE implements 3 new image classes to facilitate analysis of the images and only loading the necessary part of the image into memory rather than the whole image, since the high resolution images from smFISH-based data can be very large.

Instead of the scale factor, which is not readily available for non-Visium data, and to keep images and geometries aligned after affine transformation and cropping, the scale factor in SPE’s imgData is not typically used by SFE’s image classes. Instead, any image class to be used in SFE must have a spatial extent, i.e. a bounding box of the minimum and maximum x and y coordinates in the unit of interest. This bounding box places the images and geometries in the same coordinate system to keep them aligned through the transformations. This requirement is enforced through the AlignedSpatialImage virtual class, which inherits from VirtualSpatialImage in SPE.

SpatRasterImage is a trivial wrapper around terra's SpatRaster S4 class by inheriting from AlignedSpatialImage to fulfill SPE's image class requirement. All SpatRaster methods from terra can be applied, including finding the spatial extent, cropping, flipping, extracting values from the raster image with a vector geometry, and subsampling. Subsampling is very important as terra's implementation is very efficient in subsampling an image of over 20 GB for plotting without loading the entire image into memory. When the image doesn't fit into memory, terra only has a pointer in R, not loading the image into memory until needed. When cropping, flipping, or transposing a very large image, terra can perform the operation chunk-wise and save the results to a new file on disk without loading the whole image into memory. The geospatial field has a long history of working with such large raster data which are common in remote sensing (e.g. Google Maps satellite). Using GDAL, terra can read and write a variety of raster formats, including GeoTIFF, and optionally Zarr and TileDB if the drivers are available. The spatial extent is embedded in these formats.

A downside of SpatRaster is that much of the image processing tradition can't be applied. Hence the ExtImage class is implemented, which is based on EBImage's Image class, with spatial extent in the metadata. All image processing tools in EBImage can be applied, such as Otsu thresholding, morphological operations, and watershed segmentation. However, the whole image is loaded into memory. SpatRasterImage can be converted to EBImage; if the image is very large, it can be subsampled with the maximum number of pixels allowed before conversion.

In Xenium, the images are in OME-TIFF format. Xenium images can't be read with terra due to a compression scheme not recognized by terra, and are often too large to fully load into memory. In addition, Xenium OME-TIFF images have a pyramid of different resolutions. These images can be read into R with the RBioFormats package which gives an R interface to the BioFormats Java package[14], including selectively reading a subset of resolutions from the pyramid or a subset of pixels. SFE implements BioFormatsImage to work with these images, which is somewhat similar to the stars class in the stars package[11], which is another main R package for raster geospatial data. Only the metadata is in memory, including the coordinates of the top left corner of the image, the spatial extent, whether the spatial extent is for the full image, and affine transformation information. The top left corner is recorded so it's possible to determine the pixel range from the original image to read after the image has been cropped or translated. When cropping and affine transformation is performed, only the metadata is changed. Affine transformation information is stored either as a name and relevant parameter value (e.g. rotation and angle of rotation) or as a linear transformation matrix and a translation vector. When multiple affine transformations are performed to the same image, the equivalent transformation matrix and translation vector are stored in the metadata. If the image has been cropped, the new spatial extent is translated into pixel ranges in the resolution of interest to be read with RBioFormats. At present, BioFormatsImage functionalities have only been tested on Xenium OME-TIFF, but we may expand them to other BioFormats in the future.

SpatRasterImage and ExtImage are interconvertible. BioFormatsImage can be loaded into memory by converting to ExtImage but a resolution must be specified. Other image classes for SFE can be implemented by inheriting from AlignedSpatialImage.

#### Read functions

Read functions for Visium, MERFISH, CosMX, Xenium, and Visium HD were developed using example datasets from the companies' websites. These datasets are outputs from the standard preprocessing software from the companies, and different versions of the software give somewhat different output structures, which have been taken into account in the read functions. For MERFISH, unpublished data from different versions of the preprocessing software were used to develop the read function.

The standard output typically has cell and nucleus polygon vertex coordinates and transcript spot coordinates in either a CSV or Parquet file or both. To speed up read and operations in memory, the fread function of data.table[15] is used to read the CSV files into R as data frames. Parquet files are read with the arrow R package as data frames. Then the coordinates are converted into simple features to allow geometric operations and predicates with sfheaders. The simple features are saved to disk as GeoParquet files for faster reading the next time the data is read from the same directory. With GeoParquet and GDAL's SQL, transcript spots can be selectively read for specific genes for visualization purposes, without loading spots of all genes into memory.

#### Operations

##### Finding bounding boxes

Bounding boxes of geometries can be found with st_bbox() in sf, and those of images (aka extents) can be found with ext(), with methods for all image classes in SFE. The result is a numeric vector with names xmin, xmax, ymin, and ymax, which may be in any order. Bounding boxes for the whole SFE object is the bounding box of the union of all geometries, and images can be optionally included. When the SFE object has multiple samples, the bounding box is found for each sample separately and a matrix is returned, whose columns correspond to samples.

##### Affine transformations

Affine transformations can be applied to either the whole SFE object (Supplementary Table 2). The transform is applied to each geometry and image, using the appropriate method for each class. To keep the geometries and images aligned, for named transformations (rotate, mirror, transpose), the center of the bounding box of the whole SFE object including images is held fixed, which means that the transformation to individual geometries and images may be different from if the transformation is applied to them separately by a translation vector.

Affine transformation methods have been implemented for each of the new image classes in SFE. For SpatRaster, terra functions are used for mirroring (flip), transposing, and translation, but it must be converted into ExtImage for rotation (terra's rotation() does something very different) and general affine transformation. For ExtImage, EBImage functions are used for mirroring, rotation, and general affine transformation. When transposing, the image array is transposed and the spatial extent changed. When translating, the spatial extent is translated without changing the image itself. All affine transformations are stored in the metadata of

BioFormatsImage without changing the original image; when read into memory as ExtImage, the stored transformation information is applied on the fly. For all 3 classes, scaling the image simply scales the spatial extent.

For geometries, the affine transformations are applied to the coordinates from st_coordinates() and then the coordinates are converted back into simple features with the sfheaders package. This is much faster than calling sf's overloaded binary operators for affine transformation.

##### Aggregation

Geometries can be used to aggregate SFE data at the cell or transcript spot level. For example, a pseudo-Visium dataset can be made by aggregating Xenium or MERFISH data using a Visium spot polygon geometry. At the cell level, a spatial predicate (defaults to st_intersects but there're others such as st_covered_by, Supplementary Figure 2) can be used, so those cells returning TRUE to each Visium spot in this predicate will be aggregated into that spot. A summary function (such as sum or mean but can be other functions) can be specified to aggregate the numeric data associated with each cell. For logical columns colData, the data is converted to numeric and summarized, so sum means the number of cells with TRUE, and mean means the proportion of cells with TRUE. Categorical columns in colData such as cell type labels are converted into list columns, each element of which is a character vector, such as of cell types of all cells assigned to the Visium spot. For sum and mean, matrix multiplication is used to speed up computation, but for other summary functions, the rows of the gene count matrix are looped through but this can be parallelized.

When aggregating at the transcript spot level, the transcript coordinates can be either directly read from disk or from rowGeometries of an SFE object. When read from disk, Apache Arrow[16] is used to read coordinates of one gene at a time without loading the whole file into memory. The coordinates are converted into POINT geometry and st_intersects is used to assign the points to the geometry of interest. When aggregating from an SFE object, the rowGeometry is converted into POINT first.

The geometry used to aggregate data can be any geometry, including but not limited to Visium spot polygons, square or hexagonal grids, cell segmentations, and nucleus segmentations.

##### Splitting

One SFE object can be split with sample_id or a spatial predicate. For the latter, for example, cells intersecting with (or covered by) different polygons for different pieces of tissue are assigned to different new SFE objects. Since SFE inherits from SCE, SCE's splitting method applies.

##### Cropping

SFE objects can be cropped with any geometry of any shape, which may or may not be a bounding box. A spatial operation (by default st_intersection) is used to obtain the geometries that remain after cropping. For example, when a bounding box is used to crop an SFE object, the resulting cell segmentation geometries will be the intersection of the cells with the bounding box. Note that this can change the geometry type of the geometries. For example, when a cell segmentation POLYGON intersects with the bounding box by a point or a line, then it will become type POINT or LINESTRING respectively instead and the resulting sf data frame for all the cell segmentations will have type GEOMETRY instead of POLYGON. Other operations, such as st_difference, can also be used. With st_difference, “cropping” removes the area specified by a geometry. Alternatively, a spatial predicate can be used to keep all the cell polygons whole. By default it is st_intersects, but it can also be st_covered_by, st_disjoint, and so on. After the geometries are used to determine which cells to keep, the assays and metadata are subsetted with the SCE method.

##### Subsetting

Splitting and cropping are special cases of subsetting. Spatial graphs and annotGeometries make subsetting SFE more complicated than subsetting SCE using row and column indices. When a subset of cells in a sample are removed, the spatial graph is either directly subsetted or reconstructed with the information stored when the graph was initially created, determined by an option SFE_graph_subset. Directly subsetting can create singletons, i.e. cells without neighbors, when there was originally none. The images of this sample are cropped to the bounding box of the remaining geometries of the sample. When all cells from a sample were removed, then its corresponding spatial graphs, annotGeometries, rowGeometries, and images are also removed.

#### Implementation of alabaster.sfe

The field of spatial transcriptomics uses many programming languages, but mostly R and Python[17]. To appeal to both communities, the Bioconductor project – originally using R – is developing Python implementations of data structures and basic data analysis functionalities[18]. The Artifact DB project at Genentech aims to develop packages for language-agnostic on-disk serialization of various data structures to facilitate multi-lingual and interoperable data analysis, including using Bioconductor in both R and Python. The suite of packages to serialize and read R objects are called alabaster[19], while the Python equivalents are called dolomite. In addition, as mentioned above, SFE objects can have on-disk components such as DelayedArray assays and SpatRaster and BioFormats images that will break when the SFE object is serialized with saveRDS() and shared. In order to make the on-disk components portable and to facilitate interoperability, we have implemented alabaster.sfe for language-agnostic on-disk serialization of SFE objects from R. dolomite-sfe will be part of a longer effort for a Python implementation of SFE.

The SPE core of the SFE object (Figure 1C) is serialized with the existing alabaster.spatial package[20], but new methods were written to save and read the new image classes. Images that are in memory are saved as TIFF files and metadata such as spatial extent are saved in the JSON file that indicates object type. SpatRaster images are saved as GeoTIFF, which includes the spatial extent in the file. Images that are out of memory are copied from the original location, so the serialized bundle can be self-contained and portable.

The vector geometries are saved as GeoParquet files with the sfarrow package, as the columnar Parquet format has smaller files and is faster to read. With the packages arrow and dplyr, the Parquet file can be efficiently queried without being entirely loaded into memory. The Parquet format is supported in many different languages including R and Python, making it interoperable.

Spatial neighborhood graphs are stored as listw objects in memory in the SFE object. They are converted into CSR sparse matrices before they are written to disk with the alabaster.matrix package.

#### Methods in examples shown in figures

Panels of Figure 1, Figure 2, and Supplementary Figure 1 are assembled with LibreOffice 24.8 Draw. The satellite image in Figure 1A is taken from Google Maps and contains parts of Inwood and Spuyten Duyvil in New York City. All analyses were performed on an Asus ROG Zephyrus laptop with AMD Ryzen™ 9 5900HS with 16 CPU cores and 40 GB of RAM, running Ubuntu 24.04 LTS, R 4.5.0, and Bioconductor 3.21. All spatial tissue plots were made with Voyager 1.9.2 unless otherwise specified. Other package versions used to make figures: SpatialFeatureExperiment 1.9.7, SFEData 1.9.1, sf 1.0-19 (GEOS 3.13.0, GDAL 3.10.0, PROJ 9.5.1), dplyr 1.1.4, ggplot2 3.5.1, tibble 3.2.1, purrr 1.0.2, scuttle 1.17.0, alabaster.sfe 0.99.5, EBImage 4.49.0, terra 1.8-15, DelayedMatrixStats 1.29.0, DelayedArray 0.33.3, bluster 1.17.0, BiocNeighbors 2.1.2, sfarrow 0.4.1, patchwork 1.3.0, and scico 1.5.0.

The mouse olfactory bulb Visium data shown in Figure 2 was downloaded from the 10X website: https://www.10xgenomics.com/datasets/adult-mouse-olfactory-bulb-1-standard-1, then read into R with read10xVisiumSFE() in SFE. All smFISH-based examples in the figures come from a subset of a human pancreatic cancer Xenium dataset from the 10X website: https://cf.10xgenomics.com/samples/xenium/2.0.0/Xenium_V1_human_Pancreas_FFPE/Xenium_V1_human_Pancreas_FFPE_outs.zip. This subset is available in the SFEData package, via the function XeniumOutput("v2"). The mouse skeletal muscle muscle dataset is from day 2 post notexin injury, from reference [21], and is available in the SFEData package. Myofiber segmentation was performed manually with the Labkit plugin of FIJI and the polygons were simplified with rmapshaper. The simplified polygons are plotted behind Visium spots in Figure 2D. The Visium and Xenium data in Figure 2C come from serial sections of lung adenocarcinoma [22], patient TSU-21 in the study. The scatterplots in Figures 2E and 2G were made with plotColData() in scater 1.35.0.

##### Image registration in Figure 2C

The fluorescent image from Xenium shows different features from the Visium H&E and the latter has some histological artifact less apparent in Xenium, making image registration challenging. A quick and dirty registration was performed by first finding the tissue outline and then aligning the tissue outlines from Visium and Xenium, matching the oddly-shaped parts of the tissue. First, both the Xenium and Visium data were read into R as SFE objects with coordinates in microns. Then the images are converted to ExtImage. Contrast Limited Adaptive Histogram Equalization[23] was used to enhance contrast in the fluorescent image from Xenium (morphology_focus, resolution 4). Then both the Visium H&E (hires) and Xenium fluorescent images are normalized to have values between 0 and 1, and otsu thresholding was applied to find the tissue masks. For Xenium, a box brush with diameter 7 pixels was used in a closing operation as the mask is very porous. Then for both the Visium and Xenium masks, holes were filled and only the largest piece was kept to remove Visium fiducials and debris. The terra package was used to convert the masks into polygons, which were then simplified with sf::st_simplify() with a distance tolerance of 50 microns; the resulting polygons have more vertices at the curvier and oddly-shaped parts of the tissue outline. Point cloud registration implemented in the LOMAR R package (version 0.5.0)[24] was used to find a transformation matrix and translation vector to register the Visium tissue outline to the Xenium one. Then this matrix and vector were applied to the entire Visium SFE object with the affine() function, transforming both the geometries and images to match the Xenium spatial coordinates. The transformed Visium spot polygons can later be used to aggregate Xenium transcript spot data to form a pseudo-Visium dataset to supplement the original true Visium data that is sparser due to lower capture efficiency.

BigWarp in FIJI can be used to manually annotate landmarks for both non-linear and affine image registration. However, a challenge is that the affine transform matrix and vector include scaling from pixel space in Visium to pixel space in Xenium, which necessitates another scaling step from the Visium hires image to the coordinate unit of the Visium spots. That the images have spatial extents and are both in micron space in SFE removes this step. Furthermore, exporting the affine transformation from BigWarp is not straightforward. Novel methods such as paste2[25] can register spatial transcriptomics serial sections with gene expression, but these methods are computationally intensive and the transformation can't be simultaneously applied to the images and annotation geometries, which can be useful for visualization.

#### Extracting raster values

In Figure 2F-G, terra::extract() was used to extract pixel values in the Visium hires image behind each Visium spot polygon. Average values across pixels and channels were used. In Figure 2G, mclust 6.1.1 was used to find the 4 clusters with total transcript count per spot and average pixel values; the smallest cluster has low total counts and high pixel values (closer to the brightfield background), spots at the edge of the tissue, most likely only partially overlapping with the tissue. In the SFE package, vector geometries are frequently used to extract raster values in unit tests to ensure that the geometries and images are properly aligned.

#### Spatial aggregation and splitting

The hexagonal grids used to demonstrate spatial aggregation in Figure 2H-I were created with sf::st_make_grid(), and Figure 2H-I were directly made with geom_sf() in ggplot2 rather than with Voyager. In Figure 2I, cell centroids and st_intersects() were used to assign cells to bins, and the data from the cells were summed in each bin. Figures 2H and J demonstrating aggregating used a smaller subset of the Xenium data cropped from the version in SFEData in order to make the transcript spots discernible in the plots.

In Figure 2K, the tissue boundary polygons were manually drawn in QuPath. The different pieces here are merely separated by a blood vessel that intersects with this tissue section; in the original 3D tissue the two sides of the blood vessel are in the same piece.

#### Spatial analysis examples

Log transformation was performed on the SFEData Xenium subset with scran::logNormCounts(), but with cell area instead of total counts as the size factor. Cells with fewer than 4 total counts were removed, and the negative control features were removed as well.

For Supplementary Figure 1B, non-spatial PCA was performed on the Xenium subset from SFEData with scater, and MULTISPATI PCA was performed with Voyager. The spatial neighborhood graph used for MULTISPATI PCA was a k-nearest-neighbor graph with k=5, row normalized. The dashed horizontal lines are the theoretical upper and lower bounds of Moran's I given the spatial neighborhood graph, computed with the moranBounds() function in Voyager. The bounds help with interpretation of Moran's I values.

For Supplementary Figure 2C, Leiden clustering was performed with bluster and igraph on the log transformed gene expression values, using all genes. Concordex clustering was performed with the concordexR package, which essentially clusters the proportion of spatial neighbors in each Leiden cluster. This way concordex identifies spatial domains. Here k-nearest-neighbor with k=30 was used to find the proportions, and k-means with k=4 was used to cluster the proportions. Local Moran's I was computed for all genes with Voyager (which uses spdep's implementation). Then Leiden clustering was applied to local Moran's I values of all genes. As local Moran's I computes a mean of neighboring values weighted by the spatial neighborhood graph edge weights, it's somewhat similar to concordex. The clusters from local Moran's I classify cells according to their neighborhoods much like in concordex.

#### Spatial operation examples

In Supplementary Figure 2, commonly used examples of geometric operations implemented in sf are demonstrated on biologically relevant data. For st_buffer(), the grid of centroids were generated with st_make_grid() before st_buffer() was applied. The st_simplify() example uses the same Xenium dataset in[22]. st_concave_hull() was applied to cell segmentation polygons in the SFEData Xenium subset, as a way to find tissue boundary. In the third row, the segmentation polygon of one randomly chosen cell from the SFEData Xenium subset was used. In the last 2 rows, the circle is one randomly chosen Visium spot from the mouse skeletal muscle dataset, and the polygons are myofibers.

#### Supplementary Table 1: Comparison of functionalities of SFE and similar packages

| | *Seurat/SeuratObject* | *Giotto* | *Staffli (Semla)* | *SpatialData* | *SparialExperiment* | *MoleculeExperiment* | *SpatialFeatureExperiment* |
|---|---|---|---|---|---|---|---|
| Repo | CRAN | None | None | pip, conda | Bioconductor | Bioconductor | Bioconductor |
| Geometry representation | Centroids, polygons, transcript spots | Centroids, polygons, transcript spots | Centroids | Centroids, polygons, transcript spots | Centroids | Transcript spots | Centroids, polygons, transcript spots |
| Spatial cropping | Yes | Yes (image only) | Yes (image only) | No | No | No | Yes |
| Affine transformation | No | No | Yes | Yes | No | No | Yes |
| Spatial aggregation | No | Yes | No | Yes | No | No | Yes |
| Raster support | Visualization | Yes | Visualization | Zarr | Visualization | No | Yes |
| Read functions | Visium, VisiumHD, Xenium, MERFISH, CosMX | Visium, Xenium, MERFISH, CosMX | Visium | Visium, VisiumHD, Xenium, MERFISH, CosMX | Visium | Xenium, MERFISH, CosMX | Visium, VisiumHD, Xenium, MERFISH, CosMX |
| Direct spatial aggregation of transcript spots | No | No | Yes | Yes | No | No | Yes |
| Spatial predicates | No | No | Yes | Yes | No | No | Yes |

Supplementary Table 2: Geometries in SFE and data structures underlying similar EDA frameworks

| Functionality | SFE | SpatialData | Seurat | Giotto | Staffli[1] | Molecule-Experiment |
|---|---|---|---|---|---|---|
| Cell/spot centroid coordinates | ✅ | ✅ | ✅ | ✅ | ✅ | |
| colGeometries | ✅ | ✅ | ✅[2] | ✅ | | |
| rowGeometries | ✅ | ✅ | | ✅ | | ✅ |
| annotGeometries | ✅ | ✅ | | | | |
| Images | ✅ | ✅ | ✅ | ✅ | ✅ | |
| Geometric operations | **Functionalities of sf:** Geometric predicates: Intersect, disjoint, covered by, touch, etc. Geometric operations: find intersection,find excluded regions, union, find area, find distance, buffer, find bounding box, spatial joins, simplify geometries, etc. **Functionalities of terra:** Extraction of raster values with vector geometries, convert raster masks to polygons. **SFE methods:** See Supp. Table 3 | Query with polygon or bbox, rasterize, find bbox, find centroid, affine transformations, spatial aggregation | | Match transcript spots to cell segmentation mask or polygon, simplify polygons, subset data with bounding box, convert raster masks to polygons | | |

[1] Not equivalent to SpatialExperiment. Staffli is a class to hold Visium image and coordinate information, used in conjunction with Seurat objects in the semla package.
[2] Only works for polygons, which suffices in most cases, while any type of geometry can in theory be used in SFE.

#### Supplementary Table 3. Geometric operations in SFE

| Category | Function | Description |
|---|---|---|
| Affine transformations | transpose() | Switch the x and y coordinates |
| | mirror() | Flip vertically or horizontally |
| | rotate() | Rotate by any degree |
| | translate() | Move in the x-y plane |
| | scale() | Scale the coordinates |
| | affine() | General affine transformation specified by a matrix and a translation vector |
| | removeEmptySpace() | Translate the coordinates so the lower left corner of the bounding box is at the origin |
| Spatial relations | aggregate() | Aggregate cell-level data by any function (e.g. sum, mean, median) according to a predicate relating a colGeometry to another geometry used in binning (e.g. st_intersects, st_covered_by). |
| | aggregateTx(), aggregateTxTech() | Create SFE object whose gene count matrix comes from counting the number of transcript spots of each gene that falls into each spatial bin. |
| | annotOp() | Apply geometric operation between an annotGeometry and colGeometry, such as to get the polygons of intersections between each Visium spot and myofibers. |
| | annotPred(), annotNPred() | Apply spatial predicate between an annotGeometry and colGeometry, such as to find whether a Visium spot is in tissue, and how many nuclei are covered by each Visium spot. |
| | annotSummary() | Summarize characteristics of annotGeometries that spatially relate to colGeometries, such as the mean area of myofibers that intersect each Visium spot. |
| | bbox() | Find the bounding box of the SFE object, including all geometries and optionally images. |
| | crop() | Crop an SFE object with any arbitrary geometry, which does not have to be box-shaped. |
| | splitByCol(), splitSamples(), splitContiguity() | Split the SFE object with a geometry, e.g. cells in different histological regions get assigned to different SFE objects. Or split by sample_id. |

#### Supplementary Table 4: Methods to find spatial neighborhood graphs in SFE and similar EDA frameworks

To find spatial graphs in the histological space for neighborhood view spatial analyses, rather than gene expression or PCA space as in graph-based clustering.

| Method | SFE | squidpy | Seurat | Giotto | STUtility/ semla |
|---|---|---|---|---|---|
| Visium spot adjacency | ✅ | ✅ | | ✅ | |
| Delaunay triangulation | ✅ | ✅ | | ✅ | |
| K nearest neighbors | ✅ | ✅ | | ✅ | ✅ |
| Distance based neighbors | ✅ | ✅ | | | |
| Gabriel (prune triangulation) [26] | ✅ | | | | |
| Relative neighbors (prune triangulation) [27] | ✅ | | | | |
| Sphere of influence (prune triangulation) [28] | ✅ | | | | |
| Polygon contiguity | ✅ | | | | |

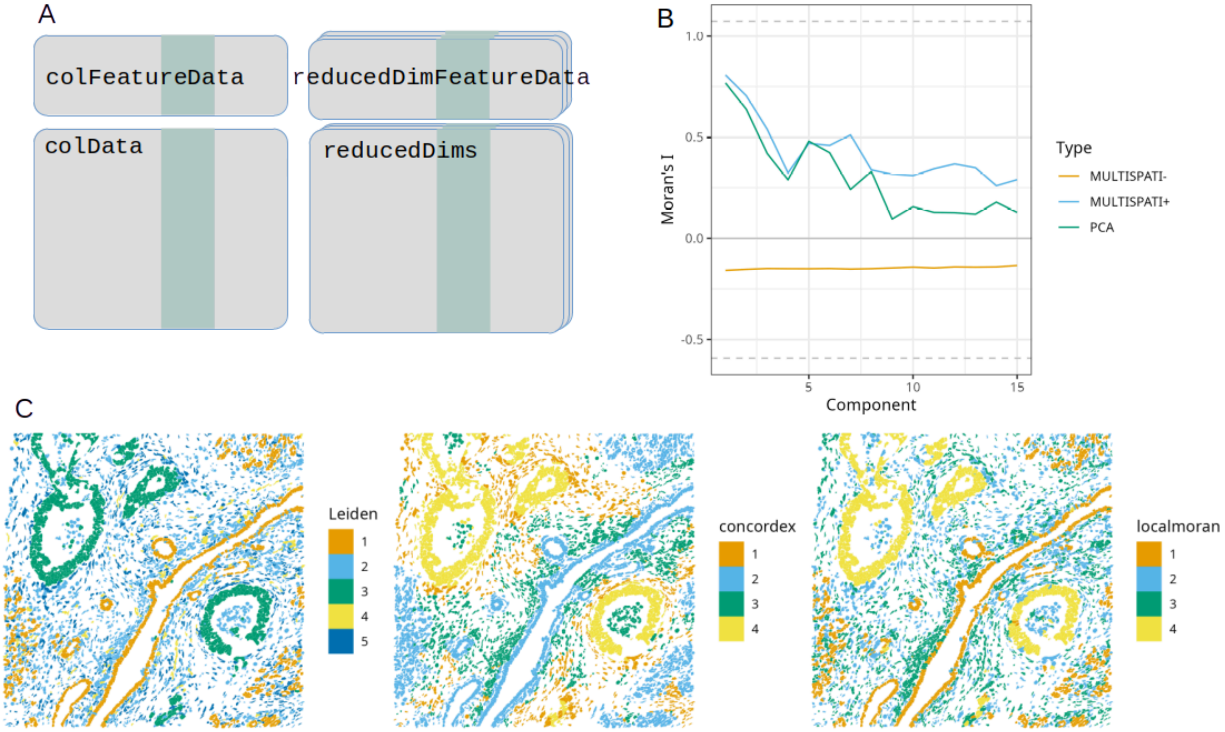


Supplementary Figure 1: A) Schematic of colFeatureData which stores global spatial analysis results for columns in colData, and reducedDimFeatureData, which stores such results for cell embeddings in each dimension reduction component such as principal components. B) Example use case of reducedDimFeatureData, plotting Moran's I of cell projections in each principal component from MULTISPATI and non-spatial PCA. The dashed horizontal lines are upper and lower bounds of Moran's I given the spatial neighborhood graph. C) Clusters identified with Leiden, concordex, and local Moran's I plotted in space.

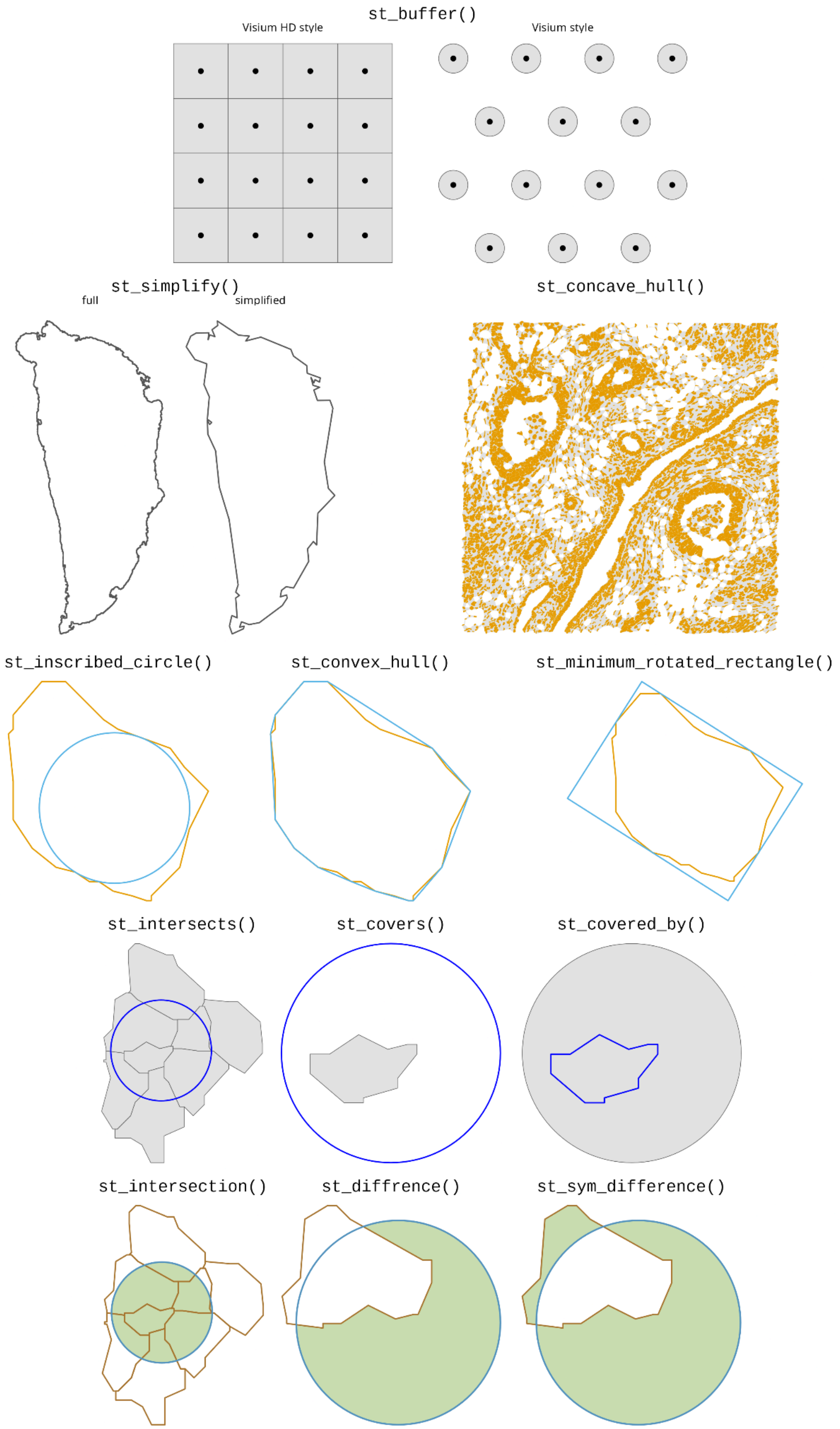
st_buffer()
Visium HD style
Visium style
st_simplify()
full
simplified
st_concave_hull()
st_inscribed_circle()
st_convex_hull()
st_minimum_rotated_rectangle()
st_intersects()
st_covers()
st_covered_by()
st_intersection()
st_diffrence()
st_sym_difference()

Supplementary Figure 2: Schematics showing examples of geometric operations in sf applied to biological data.
